# How to pour a cup of coffee

**DOI:** 10.64898/2026.08.26.746627

**Authors:** Niteesh Midlagajni, Roland W. Fleming, Constantin A. Rothkopf

**Author notes:** Correspondence and requests for materials should be addressed to Constantin Rothkopf. Correspondence.

## Abstract

Pouring a drink feels deceptively trivial, yet it requires guiding a boundary-free fluid into a vessel without spilling, overflowing, or toppling it — a task at which robots remain notoriously brittle. How humans achieve this so effortlessly is unknown, as motor control has predominantly been studied in brief, highly constrained laboratory tasks, leaving the control principles underlying ecological tasks largely unknown. Here we measured continuous sensorimotor control during liquid pouring across various containers, vessels, and speed demands. Despite substantial variation in movement trajectories and durations, individuals maintained a strikingly invariant preferred fill level. Counterintuitively, fill level variability decreased at higher fill levels, and precision was maintained even under time pressure. A stochastic optimal control model combining a data-driven nonlinear approximation of flow dynamics with a cost that balanced individualised fill level, energy expenditure and flow-rate reproduced the behaviour. Humans thus pour optimally, given their sensorimotor limits and idiosyncratic notion of “full”.

## Introduction

One of the great ironies of the human condition is that our very competence in certain domains hides from us the extraordinary complexity of the tasks we excel at. The dexterity, ease and robustness with which humans perform everyday tasks like shopping, cooking or serving a glass of wine belies how challenging they actually are. This idea is encapsulated in Moravec’s paradox (Moravec, 1988), which asserts that the tasks humans find easiest are those that are hardest to manifest in engineered systems like computers and robots. Indeed, while superhuman chess solvers emerged decades ago, there are still no robots with the sensorimotor skills of a three-year-old child, who can pluck raspberries or cut paper into shapes with scissors and robot pouring remains an open challenge in ‘physical artificial intelligence’.

How can we make headway on understanding such competences? We reasoned that by combining detailed measurements of natural human behaviour with interpretable computational models, we can infer the underlying cost functions, sources of uncertainty and control processes that enable such reliable and robust performance. Here we focused on liquid pouring. While it might seem mundane, pouring a cup of coffee is actually an exceptionally challenging control problem with numerous failure modes (Fig. 1).

**Figure 1.**
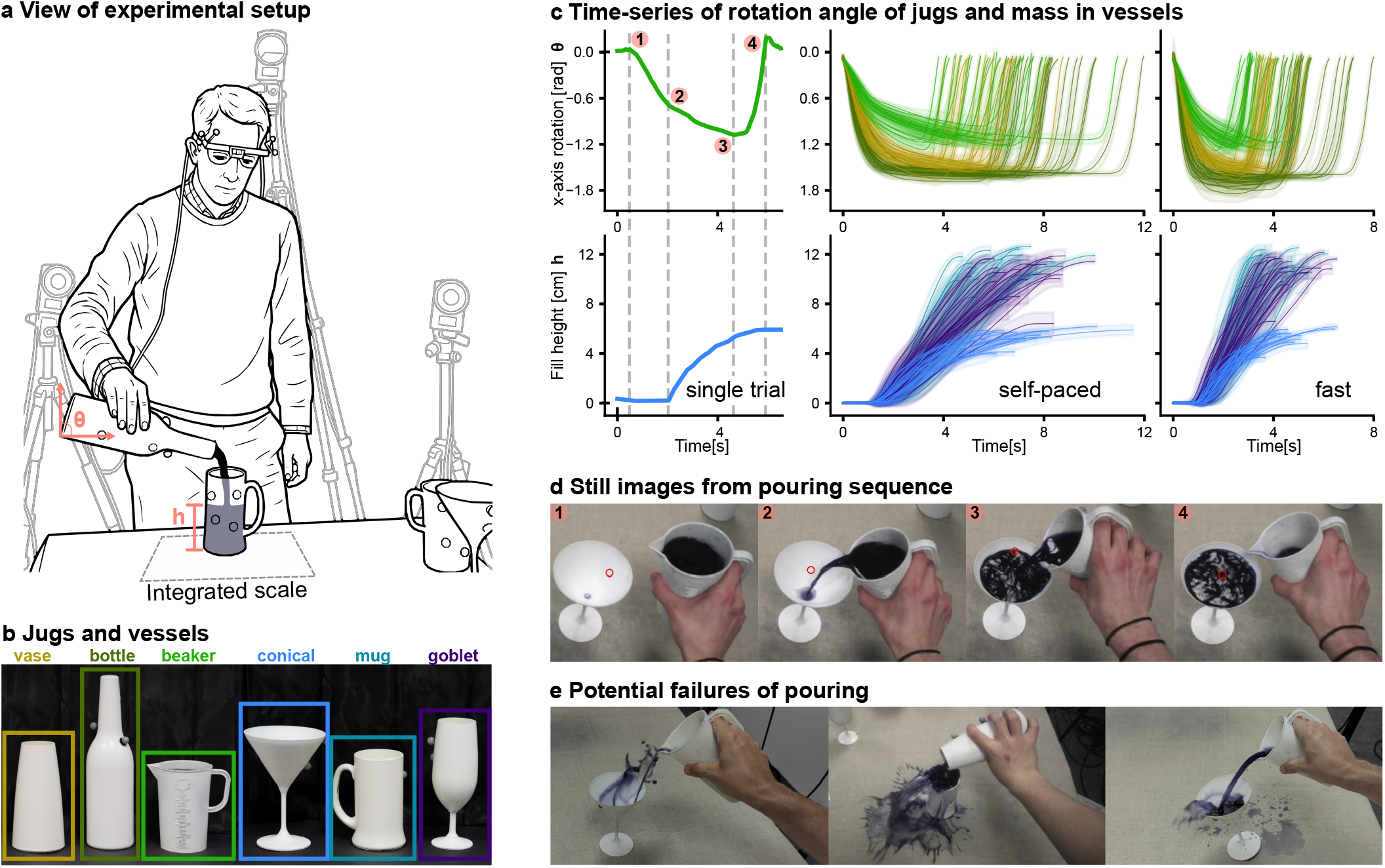
Experimental setup and behavioural data. (a) Participants’ continuous control of pouring was measured both through rotation of the respective container (via mocap), as well as the momentary amount of liquid poured, measured with a custom-built scale integrated in the table. (b) Three containers and three vessels used in the experiment. Coloured frames indicate correspondence with the traces in panel c. (c) Time series of rotational angle in radians (greens) of the three containers during a single trial (left), the self-paced (center), and the fast (right) pouring conditions, along with the fill level in cm of the three vessels (blues). Pink dots indicate four key moments in the time course: (1) initiating the rotation, (2) the fluid starts to flow, (3) reversing to slow the pour rate and (4) end of fluid flow. Note the high variability in time-course across trials, especially of the control action (container rotation angle). Despite this, we find high consistency in the outcomes of the pouring. (d) Sequence of still images from one trial from the perspective of the participant, along with the time-course of container angle (green trace) and fill level in the recipient vessel (blue trace). (e) Potential failures of pouring: spillage due to high flow-rate (left), spillage due to failure of stopping (center), failure due to toppling glass (right).

Pouring a liquid from one vessel into another is something humans generally excel at, despite the complex dynamics of liquid flow, the challenges presented by different shapes of containers and vessels—which necessitate different motor actions to achieve the same result—as well as noisy, dynamically changing environments and varying task priorities. At its heart, pouring involves rotating the source container until the fluid pours, and then rotating it back to stem the flow in time to avoid exceeding the target fill level. Yet the precise moment-to-moment time course of these rotations is highly context dependent and requires incorporating fluid flow predictions and sensory feedback while compensating for sensorimotor uncertainties and delays as the container continuously changes weight. Rotating too little can cause trickles or misses; rotating too much can topple the vessel, cause splashes or overflow (Fig. 1e). Liquid pouring is therefore a paradigmatic example of a task that humans take for granted but that even state-of-the-art robots struggle with (Reyes-Montiel et al., 2026; Rud et al., 2025). We reasoned that while human motor control in highly controlled laboratory tasks involving short, restricted actions are well studied, investigating how humans pour liquids could reveal general control principles underlying natural actions in ecological tasks.

Previous studies have described human visual perception of liquids (Kawabe et al., 2015; Paulun et al., 2015; Van Assen et al., 2018; van Assen et al., 2020), revealing sensory features that are informative about liquids’ physical properties, while others have found evidence for approximate internal models of predicting fluids’ flow (Bates et al., 2019). Others have investigated the balancing of cups filled with coffee (Mayer & Krechetnikov, 2012; Nasseroleslami et al., 2014), which is also a challenging control problem, yet inherently more stable than pouring. Seminal work on natural tasks by Michael Land (Land et al., 1999) and Mary Hayhoe (Hayhoe et al., 2003) investigated gaze behaviour when making tea or peanut-butter-and-jelly sandwiches, and both studies included pouring drinks. Yet the detailed, individual-by-individual and moment-by-moment sensorimotor control of the pouring act itself has not yet been characterised.

Here, we sought to measure and model the continuous control involved in pouring liquids from a variety of source containers into a variety of recipient vessels (Fig. 1). By design, succeeding across all conditions required distinct motor control programs for each context to account for the different pouring and filling rates. We continuously recorded the exact volume poured using a custom-built scale, while the movements of the vessels and the liquid were reconstructed from motion-capture data. To understand the underlying control processes, we turned to stochastic optimal feedback control models (Shadmehr & Mussa-Ivaldi, 2012; Todorov, 2005; Wolpert & Ghahramani, 2000). Because a closed-form description of the dynamics of flowing liquids is impractical, we used a data-driven nonlinear dynamical system approximation (Brunton et al., 2016a) to capture the flow dynamics. We find that combining the approximate dynamics with optimal feedback control not only reproduces the moment-to-moment time course of the rotation angles and fill levels but also provides an interpretable decomposition of the goals, costs and noise sources driving each participant’s behaviour. Taken together, our findings suggest that humans pour optimally given the limits of their sensorimotor systems and their idiosyncratic notion of what counts as ‘full’.

## Results

To investigate the moment-to-moment control of pouring a liquid as in everyday filling of a cup of coffee, 20 participants repeatedly poured dark-coloured water Fig. 1a from three different *containers* (bottle, beaker, and vase) into three *vessels* (conical, mug, and goblet glass) Fig. 1b. These containers and vessels were selected such that the angles of rotation of the containers would lead to different rates of liquid flow, and the vessels would fill at different rates over time, given their shapes. Participants filled each combination of container and vessel ten times under two conditions, namely *self-paced* and *fast*, resulting in a total of 180 trials. Participants were not given any instructions other than to fill the vessels as they would in everyday situations in the self-paced condition, or as quickly as they could without spilling any liquid in the fast condition. Crucially, the definition of ‘filling’ was left to the participants: obviously, a few drops would be insufficient, while filling to the very brim would be risky, but they had to decide for themselves what felt natural. The experimental setup enabled tracking of the position and rotation angles of the containers during pouring, as well as the gaze direction using an eye tracker. Additionally, the table on which the pouring task was carried out contained a custom-built scale that continuously measured the mass of the liquid in the recipient vessel. This allowed obtaining the time series of both the participants’ control of containers’ rotation angle (Fig. 1c, top), alongside its consequence, the rising liquid level, for every trial (Fig. 1c, bottom). Key moments in the pouring process are highlighted with the numbered pink dots, as explained in the figure caption. We found that the temporal profiles of both signals varied substantially across containers, participants, experimental conditions, and individual trials. Rotation trajectories differed in both the magnitude and timing of container tilts. The liquid rose at correspondingly different rates, over trials differing several-fold in duration. We therefore asked if there were consistent structures underlying these heterogeneous trajectories.

### Pouring under time pressure does not change how much people pour

We first examined how the task affects the pouring behaviour. In the fast condition, pouring duration was significantly shorter than in the self-paced condition (repeated-measures ANOVA, *F* = 117.1, *p <* 0.001), a reduction of 36.4%, and the average fill rate was correspondingly larger (*F* = 117.37, *p <* 0.001). A higher flow-rate into the same vessel means that the liquid level rises more steeply, and any error in the timing of the stopping action costs proportionally more. Yet, surprisingly, the total amount of liquid poured was maintained across conditions (*F* = 1.62, *p* = 0.22; Fig. 2a), meaning that participants hit their respective fill targets even when the dynamics of the task altered drastically. Projecting each participant’s mean fill weight onto the geometry of each vessel shows the same pattern from the perspective of the vessel (Fig. 2b): individuals differ from one another in where they stop, while each participant’s own stopping points are similar across all three vessels.

**Figure 2.**
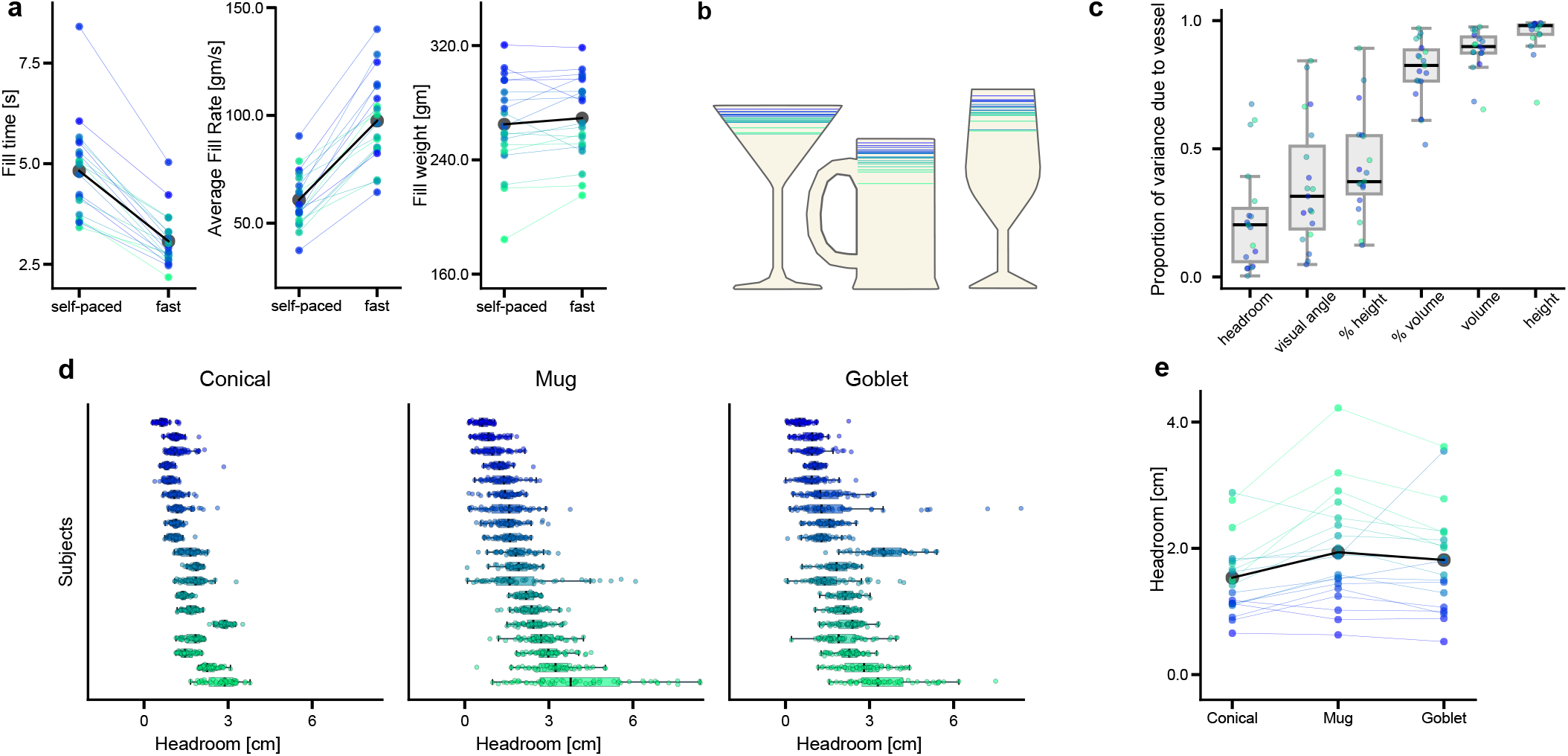
Behavioural invariants. (a) Fill time, average fill rate, and fill weight per participant across self-paced and fast conditions (individual participants in colour; group mean in black). Despite pouring faster, participants poured the same amount of liquid in both conditions. (b) Average fill weight per participant converted to the respective fill height and projected onto the corresponding vessel geometries (same colour scheme as in panel a). (c) Proportion of each participant’s variance attributable to the vessel, for six candidate stopping criteria (headroom, visual angle, % height, % volume, volume, height). Box plot shows the distribution across participants; each point is one participant. Headroom showed the lowest vessel dependence, identifying it as the quantity participants held most constant across vessels. (d) Distribution of headroom per participant across the three vessels (Conical, Mug, Goblet). Each row is one participant; points are individual trials. Note the generally consistent ordering of values across participants for the three vessels, indicating idiosyncratic preferred stopping values. (e) Mean headroom per participant across the three vessels (individual participants in colour; group mean in black). Note again the high degree of consistency within participants, especially relative to the variations between participants.

### Participants use headroom to determine when the vessel is full

To better understand the regularities in individuals’ behaviour across conditions, we next asked what physical or sensory criterion participants might use to determine when the vessel was full. We compared six plausible hypotheses about how they decided to stop pouring. These criteria spanned both a sensorimotor heuristic (Gibson, 1979; O’Regan & Noë, 2001)—specifically (1) the visual angle of the liquid surface relative to the rim—as well as criteria that involve a more structured internal representation of the pouring task (Wolpert et al., 2011; Zhu et al., 2020), specifically (2) the absolute height of the liquid, (3) the total volume poured, the proportion of the vessel’s (4) height or (5) volume filled, and (6) the distance of the liquid level from the vessel’s rim—i.e. the ‘headroom’. We reasoned that if participants actively regulate a specific measure, it should remain relatively stable across conditions and vessels, whereas a quantity that merely covaries with vessel geometry should not. We quantified this as the proportion of each participant’s total variance attributable to the vessel (see Methods), a dimensionless index that is low for a regulated measure and high for one that is not.

The results separate the candidate criteria broadly into two groups (Fig. 2c). Absolute measures of volume (median 0.899) and height (0.981), along with relative volume (0.825), are near the ceiling, indicating that participants neither execute a stereotypical container movement across vessels to achieve the same fill volume, nor adapt their movements to target the same absolute height. In contrast, spatial quantities that measure the remaining gap at the top of the vessel fared better: headroom (0.203), visual angle (0.315) and relative fill height (0.372, which captures how full a given vessel is and is thus akin to the remaining space). Among these, headroom showed the lowest vessel dependence. Bootstrapping over participants, each of the other five measures was reliably higher than headroom (relative fill height +0.19, 95 % CI [0.12, 0.36]; relative volume +0.64, [0.56, 0.76]; volume +0.70, [0.64, 0.82]; height +0.78, [0.73, 0.90]). The difference from visual angle, while positive, was small (+0.12, 95 % CI [0.02, 0.29]), and at the individual level, headroom was the best regulated measure for 12 of 19 participants, visual angle for five and relative fill height for two. This is expected given the task design: because participants were given complete freedom in their posture and viewing position in order to capture natural behaviour, they adopted a comfortable and fairly consistent stance, making headroom and visual angle effectively two parameterisations of the same physical gap. Since headroom was the quantity tracked most closely by the majority of participants, we adopt it as the measure of fill level throughout.

### Participants have idiosyncratic preferred fill levels

We next asked what determines the value of the headroom. To separate the contributions of vessel, container, task condition and participant, we fitted a Bayesian hierarchical linear model of headroom with random intercepts for participants (see Methods). The model revealed substantial between-subject variability in overall fill level (standard deviation of 0.781 cm, Highest-density interval, HDI: [0.531, 1.065]) relative to within-subject residual variability (standard deviation of 0.691 cm, HDI: [0.674, 0.707]), yielding a mean intraclass correlation coefficient (ICC) of 0.552 (HDI: [0.388, 0.715]). Thus, 55.2% of the total variance in fill level was attributable to stable individual differences.

Container type did not affect fill levels, despite each of them having unique liquid flow dynamics (Bottle: 0.044 cm, HDI: [-0.016, 0.101]; Vase: -0.034 cm, HDI: [-0.091, 0.023]; Beaker as reference). The fast condition had only a small effect, with participants leaving marginally less headroom (-0.14 cm, HDI: [-0.18, -0.09]). This difference corresponds to just 4.3 g of additional liquid, equivalent to approximately 44 ms of pouring at the fast-condition average fill rate of 97.5 g/s, and is consistent with the preserved total fill volume reported above. Vessel shape exerted a small but systematic influence: relative to the Mug, participants left less headroom in the Conical vessel (−0.42 cm, HDI [−0.48, −0.37]) and slightly less in the Goblet (−0.13 cm, HDI [−0.19, −0.07]), against a mean headroom of approximately 1.8 cm. Note that these vessel effects are small, indicating that participants remained close to their preferred fill level across vessels. This individual consistency is directly visible in the data (Fig. 2e): each participant’s mean headroom is nearly constant across the three vessels, and the ordering across participants is almost perfectly preserved, such that a participant who leaves a large gap in one vessel generally does so in all three. The underlying trial-by-trial distributions (Fig. 2d) confirm the consistency of these preferences, and reveal a second form of individual difference: participants differ not only in their fill levels, but also in how tightly their pours cluster around that point.

### Variability scales with the preferred headroom

The trial-by-trial distributions in Fig. 2d show that participants differ in how tightly their pours cluster around their preferred level. To characterise this variability, we computed, for each participant, vessel and condition, the standard deviation of the headroom across the ten repetitions and related it to the corresponding mean (Fig. 3a). Across participants, variability increased with the mean headroom; participants who filled closer to the rim were more precise across repetitions. The relationship was positive and significant in all three vessels (linear regression; Conical *β* = 0.054, *R*^2^ = 0.097, *p <* 0.001; Mug *β* = 0.200, *R*^2^ = 0.376, *p <* 0.001; Goblet *β* = 0.173, *R*^2^ = 0.313, *p <* 0.001), and held separately in both conditions. Such scaling of variability with the magnitude of the controlled quantity is characteristic of signal-dependent noise in sensorimotor control (Harris & Wolpert, 1998).

**Figure 3.**
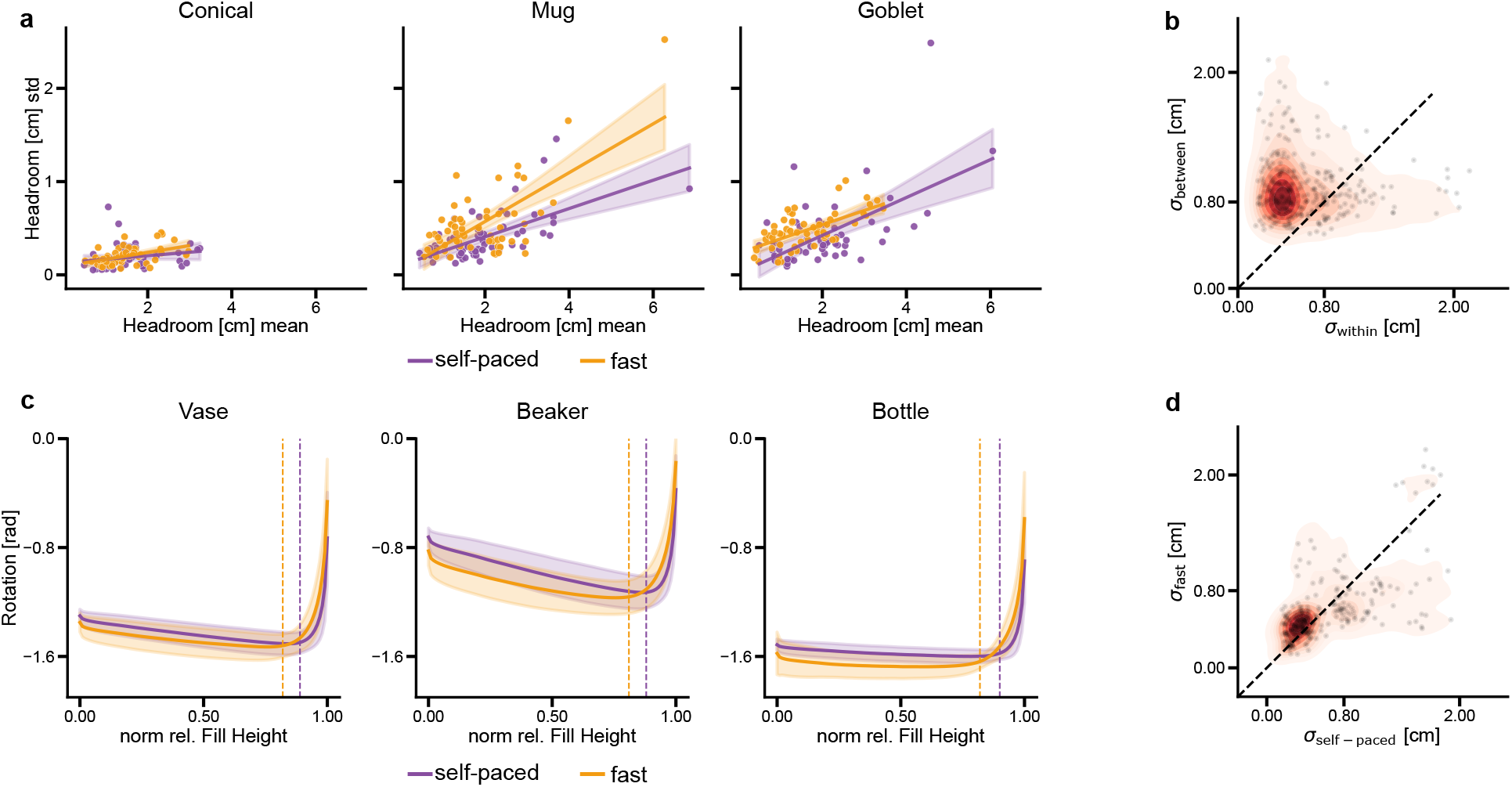
(a) Standard deviation of participants’ headroom as a function of their mean headroom, for self-paced (purple) and fast (orange) conditions across the three vessels. Participants who filled closer to the rim (lower mean headroom) exhibited lower variability. (b) Between-participant variability (*σ*_between_) exceeds within-participant variability (*σ*_within_); 300 of 20,000 bootstrap samples shown for clarity. (c) Container rotation about the pouring axis as a function of normalised fill height, averaged across participants for each container type (Vase, Beaker, Bottle). In the fast condition (orange), participants rotated the container further, increasing liquid flow, and reversed the rotation earlier to compensate for the greater residual outflow and reach the same fill level as in the self-paced condition (purple). Dotted vertical lines indicate inflection points. (d) Participants’ variability was similar in the fast (*σ*_fast_) and self-paced (*σ*_self−paced_) conditions; bootstrap analysis (300 of 20,000 samples shown) confirms that task speed did not systematically alter pouring variability.

Moreover, this variability is not solely a consequence of the fill level itself, but reflects a stable individual characteristic. A bootstrap analysis comparing variability computed within participants against variability computed across participants confirms this (Fig. 3b): drawing trials from a single participant consistently yields a smaller spread than drawing the same number of trials from the remaining participants pooled (see Methods). Thus, where a participant stops and how precisely they stop are both stable individual characteristics.

### Precision is maintained under time pressure

The ubiquitous speed-accuracy trade-off in human movement, often known as Fitts’s Law (Fitts, 1954; Woodworth, 1899), predicts that the increased speed demand of any given task should carry an accuracy cost. Accordingly, in the fast condition, a pour shortened by roughly a third (36.4%), delivering the same amount of liquid into the same vessel at a higher fill rate, should terminate at a more variable fill level. Strikingly, this was not the case. Individual participants’ variability did not differ between the self-paced and fast conditions (repeated-measures ANOVA, *F* = 1.08, *p* = 0.31; Fig. 3d). Participants reached their individually preferred headroom with the same precision in substantially less time.

The container rotation profiles reveal how participants achieved this. Plotting container rotation as a function of the normalised fill level, which aligns trials of different durations by how far the filling has progressed, shows that the control differed systematically between the fast and self-paced conditions for all three containers (Fig. 3c). First, participants rotated the container further, increasing the flow-rate, which results in vessels filling faster. Second, participants reversed the rotation earlier in the pour, with the inflection point of the rotation profile shifting to a lower fill level (dashed lines) by 6.7% of the vessel’s fill height on average. A faster pour results in more residual liquid outflow after the rotation reverses, requiring an earlier stopping movement to reach the same fill level as in the self-paced condition.

Overall, pouring behaviour was dominated by idiosyncratic preferred fill levels that individual participants maintained across trials, despite changes in the shapes and sizes of both containers and vessels, and across both task conditions. This consistency, maintained even as the dynamics of the task changed substantially, hints that participants’ moment-to-moment control and criterion for stopping are guided by a structured internal model of the physical scene, including an approximate representation of the liquid dynamics, rather than by simple sensorimotor heuristics. We tested this by implementing a model of the pouring process.

### An approximate model of fluid dynamics is sufficient to describe pouring

To better understand the observed moment-to-moment control, we turn to the framework of stochastic optimal feedback control under uncertainty (Åström, 1965), which accounts for a wide range of sensorimotor behaviours, including basic motor tasks such as pointing and reaching (Shadmehr & Mussa-Ivaldi, 2012; Todorov, 2005; Wolpert & Ghahramani, 2000), and recently to more complex behaviours in ecological tasks including ball-catching (Belousov et al., 2016) and navigation (Kessler et al., 2024). Applying this framework requires formulating the dynamical system describing the temporal evolution of task-relevant variables. However, the complex dynamics of flowing liquids are notoriously difficult to capture (Batchelor, 2000). This is one of the reasons why, despite significant advancements in robotic perception and control, achieving human-level dexterity and precision in pouring remains essentially unsolved (Reyes-Montiel et al., 2026; Rud et al., 2025). Human perceptual judgements, however, are consistent with an approximate rather than exact internal model of liquid behaviour (Bates et al., 2019; Ullman et al., 2017). We hypothesized that an approximate model relating the angle at which the container is held to the resulting flow-rate might be sufficient to perform the task, as recent research has shown that simplified internal models can account for the control of complex object manipulation (Bazzi et al., 2024).

To obtain such a model, we used sparse identification of nonlinear dynamics with control inputs (SINDYc) (Brunton et al., 2016a, 2016b), which recovers minimal differential equation models from data using a library of candidate functions. We defined the system state as the liquid weight *w*(*t*), its rate of change 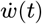, the container rotation *θ*(*t*) and the headroom *d*(*t*), driven by a single control input: the container rotation rate 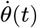. SINDYc was then used to identify, for each container, the nonlinear function governing how these state variables evolve over time (see Methods) (Fig. 4a). The model was trained on 80% of per-container pouring trajectories and evaluated on the remaining 20%. To validate the identified dynamics, we replayed participants’ measured rotational control through the learned system and compared the simulated liquid flow with the quantities actually poured. The identified dynamics reproduced the observed fill time courses on held-out trials across all containers and vessels (Fig. 4b; RMSE 15.31, 23.65 and 19.24 across all state variables for Beaker, Vase and Bottle, respectively).

**Figure 4.**
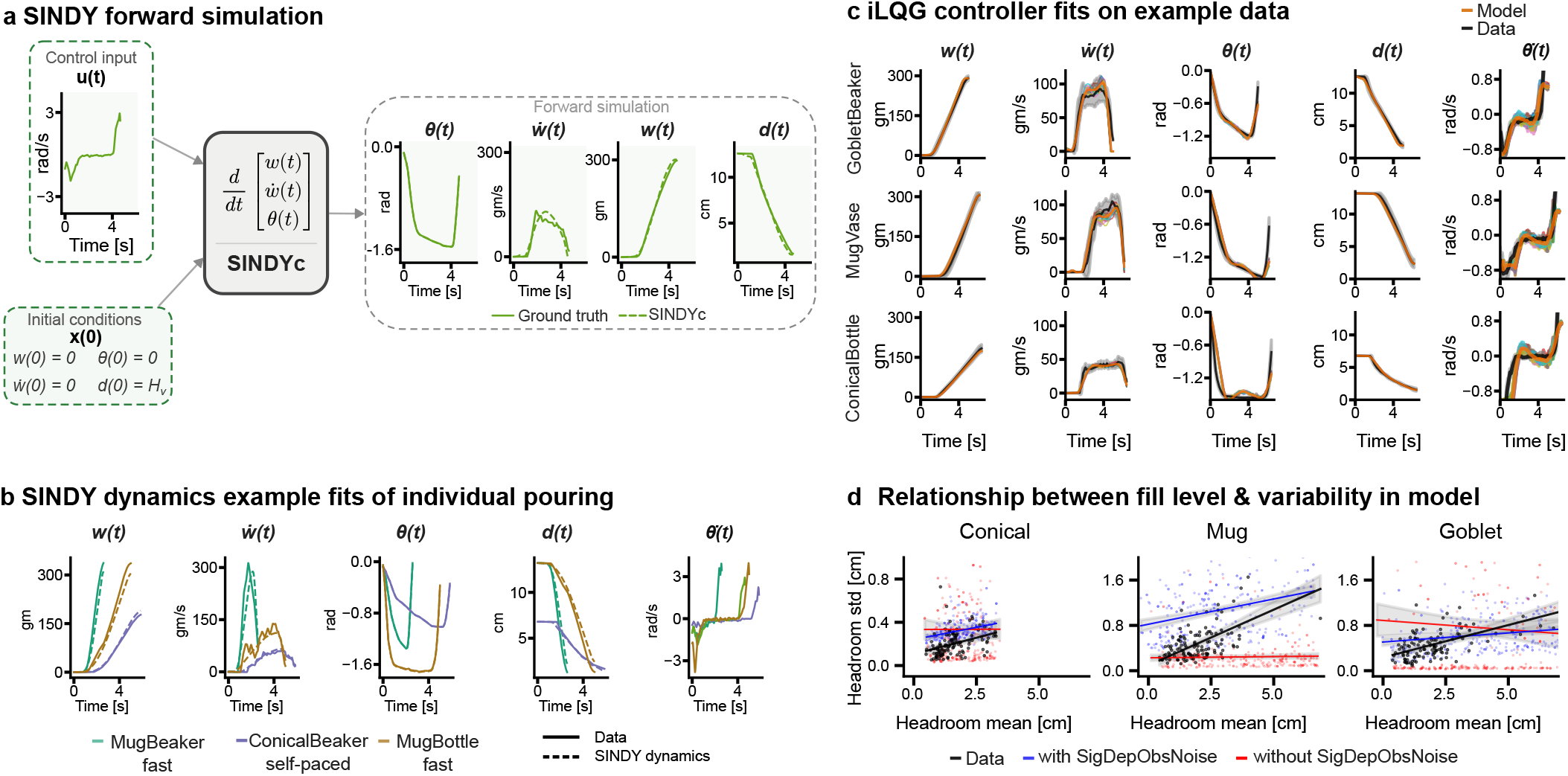
Optimal feedback control model of pouring. (a) SINDY forward simulation pipeline. (b) Example fits from SINDY forward simulation across vessels and containers showing close agreement between observed (solid) and simulated (dashed) trajectories for all state variables. Note, predictions are for test trials excluded from the training set. (c) Example tuned fits of the iLQG model (orange) to human data (black) for three Container–Vessel pairs (GobletVase, MugVase, ConicalBottle). (d) The positive relationship between headroom mean and variability observed in human data emerges from the iLQG model only when signal-dependent observation noise is included. Without signal-dependent noise, the model fails to reproduce this relationship.

### Optimal feedback control model reproduces human pouring behaviour

Given that the identified dynamical systems reproduced fluid flow, we combined them with optimal control models of human sensorimotor behaviour (Todorov, 2005). This additionally requires specifying a cost function that expresses the goals and constraints of the pouring task. We considered four components, each motivated by the motor control literature and our empirical observations. These included: (1) a control cost (Shadmehr & Mussa-Ivaldi, 2012; Shadmehr et al., 2016; Todorov, 2005; Wolpert & Ghahramani, 2000), capturing the effort of rotating the container, (2) a cost for regulating the flow-rate, capturing the task goal of avoiding spillage (Hynninen & Kyrki, 2026), (3) a cost to reach individuals’ preferred headroom *D*_tgt_, reflecting our finding that participants regulate this quantity, and (4) a cost for the container to return to its upright resting position at the end of the pour, as formalized in Equation 1:

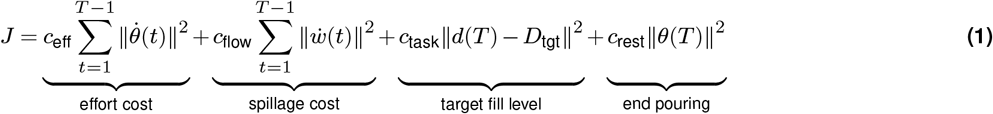

Using this cost function together with the approximate dynamical system, we computed the optimal feedback control policy assuming signal-dependent motor and observation noise (Todorov, 2005) (see Methods). By tuning the cost function parameters for representative container–vessel combinations, the model reproduces the observed moment-to-moment behaviour, including the rotation of the container and the resulting liquid variables (Fig. 4c).

Closer inspection of the model reveals the source of two behavioural phenomena reported above. First, the positive relationship between mean headroom and its variability (Fig. 3a) emerges only when simulating the model with signal-dependent observation noise that scales with headroom; the same model with additive observation noise does not reproduce it (see Methods) (Fig. 4d). Second, the model preserves the precision of the final fill level under increased pouring speed (fast condition), consistent with the empirical data. The controller adjusts the timing of the rotation reversal to compensate for the higher flow-rate, rather than executing the same movement faster. Together, this demonstrates that the identified dynamics and cost function are sufficiently expressive to capture human pouring, and motivates performing inverse modelling of the behaviour, to systematically recover the underlying parameters from individual participants’ data.

### Inverting the model recovers individual goals and costs

Having established that the optimal feedback control model captures the moment-to-moment kinematics of pouring, we leveraged the interpretability of our cost function’s parameters to invert the model to estimate individual behavioural strategies from the observed data. Specifically, we used nonlinear least-squares trajectory fitting to recover each participant’s effort cost *c*_eff_, flow-rate cost *c*_flow_, and preferred headroom *D*_tgt_ from the observed pouring data, separately for each container, vessel and condition (see Methods).

We first verified that the estimation procedure reliably identifies the underlying parameters. Mapping the objective landscape for individual participants confirmed that the optimisation reliably converges to the global minimum (Fig. 5a). We then assessed whether each component of the cost function is necessary to capture human behaviour by comparing the full model against reduced variants using ΔBIC (Bayesian Information Criterion) scores. Removing the flow-rate cost produced a measurable deterioration in fit across all container–vessel combinations and conditions, while removing the target headroom cost caused a severe deterioration, consistent with the model having no incentive to pour without a fill target (Fig. 5b).

**Figure 5.**
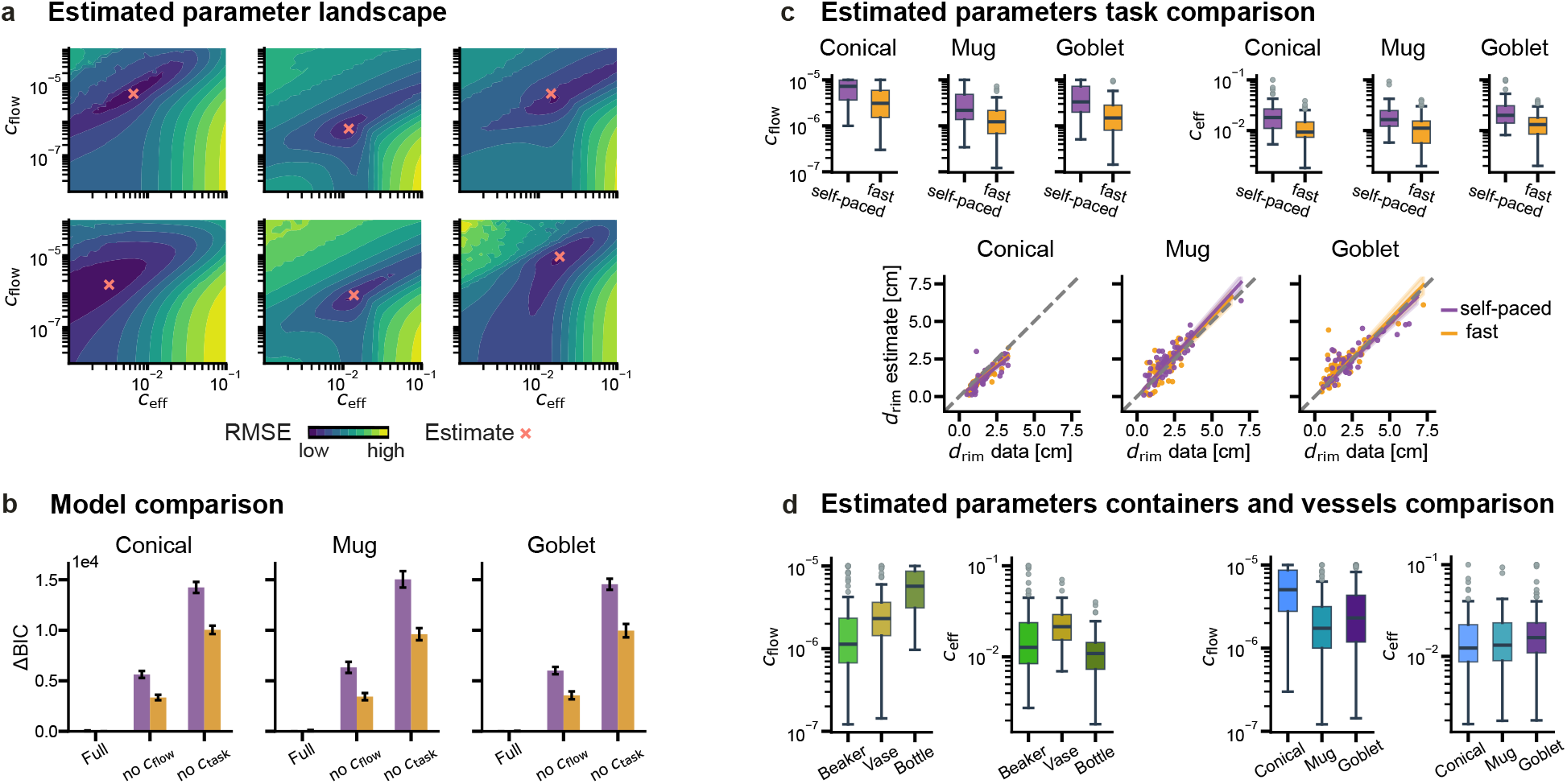
Inverse optimal control results. (a) Objective function landscape for example participants. The estimated parameters (indicated by ‘x’) lies in the vicinity of the landscape minimum, indicating reliable convergence of the optimisation. (b) Model comparison using ΔBIC between the full model and reduced variants fit to human data. Removing the fill-rate cost substantially degrades model fit across all combinations and conditions, demonstrating that regulation of fill rate is an essential component of the control strategy. Models excluding fill cost perform worst. Action cost is not shown, as optimisation fails to converge in its absence. (c) Recovery of target headroom and inferred cost parameters from human data. Recovered headroom versus measured headroom (bottom row) and inferred fill-rate cost and action cost (top row) for Conical, Mug, and Goblet pours, shown separately for self-paced (purple) and fast (orange) conditions. Both fill-rate and action costs are consistently lower in the fast condition, indicating that participants expend more energy and deregulate fill rate to achieve faster task completion. (d) Inferred costs differ systematically across both containers and vessels. When comparing across containers, the Bottle exhibits a higher fill-rate cost than the Beaker and Vase, indicating greater difficulty in regulating liquid flow. When comparing across vessels, the Conical glass shows higher fill-rate cost than the Mug and Goblet glasses, reflecting increased difficulty in achieving accurate filling without spillage. This is consistent with participants’ post-experiment difficulty ratings, which identified the Conical+Bottle combination as the most difficult.

The recovered parameters reveal several interpretable findings about the pouring behaviour and quantify them precisely. First, the estimated preferred headroom *D*_tgt_ agreed closely with each participant’s empirically observed idiosyncratic stopping criteria (Fig. 5c, bottom row), confirming that the model identifies the same goal that was inferred directly from the data. Second, comparing self-paced and fast conditions revealed a consistent shift in cost weights: under time pressure, both *c*_eff_ and *c*_flow_ were reduced across all vessels (Fig. 5c, top row). This suggests that, in order to complete the task faster while achieving the same fill target, participants were willing to expend more effort and accept a greater risk of spillage. Finally, the inferred costs quantify the computational demands that different containers and vessels impose on pouring performance. Among vessels, across all tasks and participants, the conical glass carried the highest flow-rate cost (Fig. 5d, right), reflecting the difficulty of regulating the fill rate in this vessel to control spillage. Crucially, the estimated control effort cost (*c*_eff_) remained virtually unchanged across vessel types, as, purely from a kinematic standpoint, pouring from a given container requires the same physical effort regardless of the receiving vessel’s shape. Among containers, the Bottle carried the highest flow-rate cost (Fig. 5d, left), consistent with its glugging dynamics, which make it harder to control the liquid flow and increase the risk of spillage. In contrast, the Beaker container yielded the lowest flow-rate cost, as its prominent spout made it easier to pour and inherently minimized the risk of spillage. Both orderings correspond to participants’ post-experiment difficulty rankings, in which the conical vessel and the Bottle were reported as the most difficult, and their combination as the hardest of all.

Taken together, the optimal feedback control model not only reproduces the moment-to-moment control of pouring but also recovers, for each individual participant, both the preferred fill level and the cost structure underlying their behaviour. These results suggest that human pouring can be described as optimal control under uncertainty, in which participants minimise effort and spillage risk while reaching an individually determined fill target.

## Discussion

Human motor control has been understood largely through brief, highly constrained laboratory movements, leaving open the question of how the nervous system controls everyday actions amid complex, uncertain, and nonlinear environmental dynamics. Liquid pouring provides a stringent test: humans perform it effortlessly across vessels and speeds, whereas robots remain brittle. We reasoned that working out how humans solve this task could reveal general principles of biological optimal control under uncertainty. The task design necessitated distinct motor control programs for each container–vessel pairing, to compensate for different pouring profiles and filling rates. Despite these substantial differences in the moment-to-moment kinematics of pouring, we found a striking, idiosyncratic preference for fill level in terms of the amount of space left at the top of the vessel — the headroom. Not only was this distance constant across different vessels and containers, but it was also maintained across both the self-paced and fast conditions, the latter without any measurable loss in precision.

It is interesting to speculate about potential causes for individuals’ idiosyncratic preferred fill levels. One possibility is that it reflects variations in sensory or motor noise, such that people with greater noise leave larger headroom to avoid spilling. A direct test of this would be to experimentally increase sensory uncertainty (e.g., blurry glasses) or motor variance (e.g., adding a vibration device to the arm) to test if participants choose lower fill levels. Accordingly, preferred fill levels might also be context dependent. For example, if the task involved passing the drink to another person or carrying it on a tray over uncertain terrain, lower fill levels might be preferred. Other factors might even include social norms about appropriate fill levels for drinks (e.g., British pint filled to the brim vs French wine glass filled barely one-fifth). This would, however, predict systematic variations in fill level across glasses, which we did not observe in our task. Previous work has shown that it is possible to disentangle motor costs from other factors in dynamic tasks (Raßbach et al., 2025), and future work could test whether fill level preferences adapt to changes in motor costs, context, or social factors.

The violation of the speed–accuracy trade-off is a surprising behavioural finding, given how robust this trade-off has been in motor control across a range of tasks (Fitts, 1954; Woodworth, 1899). Participants nearly halved their pouring duration without any loss of precision in the final fill level. This can sound paradoxical if we presume the same movement was executed faster. What the kinematics show instead is that the motor control was qualitatively reorganised, i.e. the container was rotated further to generate higher flow, and the stopping point was initiated at a lower fill level to absorb the larger residual outflow. The headroom was thus maintained by adapting the strategy. This is the signature of optimal feedback control organised around a task goal rather than the trajectory, in which variability is permitted along task and control dimensions that do not affect the outcome and suppressed along those that do (Todorov & Jordan, 2002).

On a broader view, the present study speaks to a resurgent interest in natural, ecological tasks both in cognitive science (Fooken et al., 2023; Goettker et al., 2025; Maselli et al., 2023; Rothkopf & Hayhoe, 2025; Schakowski et al., 2026) and neuroscience (Angelaki et al., 2026; Cisek & Green, 2024; Krakauer et al., 2017; Miller et al., 2022; Yoo et al., 2021). Yet, a central challenge in computationally modelling such behaviour in humans is extending normative frameworks, such as stochastic optimal control, beyond simple paradigms in which task dynamics are tractable and derived from first principles (Shadmehr & Mussa-Ivaldi, 2012; Todorov, 2005; Wolpert & Ghahramani, 2000). Expanding this framework to everyday ecological, natural tasks has had a few successes, including ball-catching (Belousov et al., 2016), striking a target with a whip (Krotov et al., 2022), and navigation (Kessler et al., 2024), but a major bottleneck is the impossibility of deriving tractable dynamical systems that capture complex nonlinear interactions. Liquid manipulation, for example, is governed by dynamics that are intractable in closed form (Batchelor, 2000). By combining data-driven system identification (Brunton et al., 2016a, 2016b) with optimal feedback control, we show that capturing these low-dimensional surrogate dynamics is sufficient to account for fine-grained human motor control, even in a task involving nonlinear dynamics as complex as pouring. This is consistent with growing evidence that skilled motor behaviour in complex tasks relies on simplifying the control problem through predictive strategies (Russo et al., 2025). This paves the way to understanding how humans solve goal-directed sensorimotor tasks by carefully isolating key task-relevant states, pairing them with data-driven system identification, and defining interpretable cost functions within a stochastic optimal control framework.

Beyond forward simulation, a crucial step toward understanding natural behaviour is inverse optimal control, which reveals why a particular policy is selected over countless alternative motor programs and, in the process, recovers the implicit cost functions and internal constraints that drive biological action (Rothkopf & Hayhoe, 2025). This approach has proven effective at uncovering internal objectives across diverse motor behaviours (Mombaur et al., 2010; Rothkopf & Ballard, 2013; Straub & Rothkopf, 2022). We demonstrate that inverse optimal control can also be applied to complex everyday tasks involving fluid manipulation. Inverting our controller revealed how participants flexibly negotiated competing internal goals: under time pressure, effort and flow-regulation costs were selectively relaxed to prioritize speed, accepting higher risk of spillage while hitting the same target headroom. This reflects recent evidence that speed in goal-directed movements arises from joint optimisation of metabolic effort and accuracy costs (Bruening et al., 2024). Furthermore, the inferred cost parameters naturally captured the physical difficulty of handling certain objects in the environment, such as assigning high flow-rate costs to the glugging bottle and the sensitive fill profile of the conical vessel. Importantly, such analyses are based on individual-by-individual, trial-by-trial, and moment-by-moment models rather than statistical averages of behavioural data. Applying inverse optimal control to natural tasks thus provides a quantitative window into how humans balance their internal costs to achieve task goals under real-world constraints.

Humans routinely perform tasks involving interactions with materials, tools, and environments governed by complex nonlinear dynamics. Examples include billowing a bedsheet onto a mattress, guiding slurry on a building site, controlling a digger, or tethering a horse. It seems unlikely that we have innate internal physics models for every such eventuality. Instead, it seems far more likely that we have a means to rapidly infer (and continually adapt) approximate generic dynamical models that relate motor actions to world outcomes (as measured through the senses). This capacity lies at the core of physical artificial intelligence, widely recognised as the next major frontier for building tractable world models and robotic systems for unstructured noisy environments (Sitti, 2021). Indeed, despite vast research, robotic pouring remains notoriously brittle in the general case (Reyes-Montiel et al., 2026; Rud et al., 2025), often requiring a dense array of sensors, whereas humans succeed using just two noisy eyes and a compliant yet noisy arm. This gap suggests that biological dexterity does not depend on sensory brute force, but on an efficient computational architecture. Uncovering how humans solve these everyday tasks may therefore prove essential not only for understanding biological sensorimotor control, but also for designing resilient embodied AI.

## Data and Code availability

Data and code generated as part of this publication is available at github.com/RothkopfLab/human-pouring-control.

## Author contributions

Conceptualization, N.M. and C.A.R. methodology, N.M. and C.A.R.; investigation, N.M. and C.A.R.; writing: N.M., C.A.R. and R.W.F.; data visualization: N.M., C.A.R. and R.W.F. funding acquisition, C.A.R. and R.W.F.; resources, N.M. and C.A.R.; supervision, C.A.R.

## Acknowledgements and Funding

We thank Dominik Straub for insightful discussions on optimal control framework. This study was funded by the European Research Council (project number ERC-CoG-101045783 455 “ACTOR” to C.A.R. and project number 101098225 - ERC-2022-AdG “STUFF” to R.W.F.) and the German Research Foundation under Germany’s Excellence Strategy (EXC 3066/1 “The Adaptive Mind”, Project No. 533717223). C.A.R. additionally acknowledges funding through the Simons Collaboration in Ecological Neuroscience (SFI-AN-NC-SCN-00007276). We gratefully acknowledge the computing time provided to us on the high-performance computer Lichtenberg at the NHR Centers NHR4CES at TU Darmstadt.

## Competing interests

The authors declare no competing interests.

## Additional information

Correspondence and requests for materials should be addressed to Constantin Rothkopf.

## METHODS

### Participants

Twenty participants took part in the experiment (26 ± 5 years; age range: 20 to 38; 13 females). Participants were undergraduate or graduate students recruited from the Technical University of Darmstadt who received course credit for their participation. All experimental procedures were carried out in accordance with the guidelines of the German Psychological Society and approved by the ethics committee of the Technical University of Darmstadt. Informed consent was obtained from all participants prior to carrying out the experiment. Participants were required to be right-hand dominant and to have normal or corrected-to-normal vision.

### Experimental Setup

The experimental setup consisted of six motion capture cameras (Qualisys AB, Gothenburg, Sweden) and a height-adjustable table with an integrated scale (Kern KFP 3V20M + CE HSE). Eye movements and the scene view were tracked with the Tobii Pro Glasses 2 mobile eye tracker (Tobii AB, Stockholm, Sweden), and markers were added to the side pieces of the eye glasses to track head movements with the Qualisys motion capture system. Visual markers were also added to the three filling containers and three target glasses, which were placed on the table (Fig. 1a-b). We selected containers whose rotation angles resulted in distinct liquid flow-rates, and vessels whose geometries caused liquid to fill at different rates over time. In every container and vessel combination, the filling container was filled with 500 ml of water, which was more volume than any target glass could hold. Water was dyed with a dark food colour to make it easier to detect the liquid level visually for computer vision algorithms. Synchronisation of all data streams was achieved using ROS (Quigley et al., 2009) as backend. Motion capture and scale data were stored independently with global ROS timestamps. The Tobii eye tracker recorded data on device with its own clock. To synchronize, we sent a timestamped ROS event to the Tobii recording via its API, which was logged as an external event alongside gaze data. All data streams were then aligned to a common timeline using these shared event timestamps.

### Experimental Protocol

Participants first wore the mobile eye tracker and were asked to stand in front of a table where the scale was integrated. They were free to stand at any distance from the table. Participants were instructed to slightly tilt their head during the task to ensure that the field of view of the mobile eye tracker adequately covered the scene. The height of the table was adjusted to each participant’s preference so that the task felt natural and comfortable to perform.

The task consisted of pouring liquid from containers into glasses under two conditions. In the *self-paced* condition, participants were instructed to pour at their normal pace. In the *fast* condition, participants were instructed to pour as quickly as possible. In both conditions, participants were instructed to pour to a level of their preference, as they would typically choose if serving themselves or their guests. They were additionally instructed to complete the pouring in one continuous motion, rather than stopping and resuming to add small amounts of liquid.

In both conditions, participants first selected a container and a glass. The container was filled with 500 ml of liquid, and participants poured the liquid into the glass. Once the pour was completed, participants were instructed to pour the liquid from the glass back into the container and repeat the process. Each participant performed 10 trials per glass. We notified them once 10 trials were completed. They then selected the next glass and repeated the task for another 10 trials. After all three glasses had been completed, participants selected the next container and repeated the procedure using the same set of glasses. In total, each participant completed 180 trials. The mobile eye tracker was calibrated every 30 trials, corresponding to the completion of three glasses for a given container, or earlier if the eye tracker was accidentally displaced between trials.

### Post-experiment difficulty rankings

After the experiment, we verbally asked participants to rank the containers and vessels according to how difficult they found them to use. Across all participants, the Conical vessel was unanimously reported as the most difficult to pour into, and the Mug most often reported as the least difficult. For containers, except for one participant who reported the Vase, everyone else reported the Bottle as the most difficult to pour from, while the Beaker was unanimously reported as the easiest.

### Data Filtering

Trials were segmented from the continuous data streams using synchronised motion capture and scale data. Each trial began with the container pick-up event, defined as the moment the container was lifted from the table, and concluded with the pour-stop event, defined as the moment the weight on the scale ceased to change. Trials were excluded from analysis if more than 10% of the data points were missing, if the liquid volume exceeded the total capacity of the vessel, or if the container’s rotation deviated from a single, continuous pouring motion. One participant was excluded due to behavioural inconsistencies. Following these exclusions, the final dataset consisted of 3,367 trials from 19 participants.

### Fill level measures

Several analyses relied on quantities derived from the recorded liquid weight and the vessel geometry. To convert liquid weight to the corresponding fill height, we leveraged the vessel geometries, modelling the Mug as a cylinder and the Conical vessel as an inverted frustum. For the Goblet, whose geometry has no analytical solution, we fit a rational function to a weight–height lookup table generated by manually recording height increments at 5–10 g liquid weight intervals. From the fill height *h*, we defined the headroom as the distance from the liquid surface to the rim:

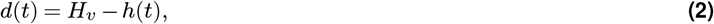

where *H*_*v*_ is the maximum fill height of vessel *v*. The headroom is thus largest when the vessel is empty and approaches zero as the liquid reaches the rim. We also expressed fill height and fill weight as proportions of each vessel’s capacity, defining the relative fill height as *h*(*t*)*/H*_*v*_ and the relative fill volume as *w*(*t*)*/W*_*v*_ , where *W*_*v*_ is the maximum liquid weight of vessel *v*.

For each trial, we additionally computed the visual angle subtended at the eye by the vessel rim and the liquid surface. Using the motion-capture data, we estimated the position of the participant’s left eye (from the eye tracker) within a 300 ms window centred on the inflection point of the container rotation, i.e. the moment at which the participant began reversing the pour. Given the known pose and geometry of the vessel, we then cast two rays from the eye position: one to the point on the vessel rim farthest from the participant, and one to the point on the boundary of the liquid surface farthest from the participant. The visual angle was defined as the angle between these two rays.

### Statistical analysis

To assess the effects of container, vessel, and task on participants’ preferred fill levels while accounting for their individual differences, we fit a Bayesian hierarchical linear model using the `bambi` Python package (Capretto et al., 2022). The model regressed headroom on vessel, container, and task as fixed effects, with a participant-level random intercept to capture between-subject variability:

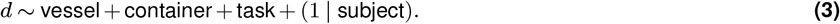

Posterior distributions over the regression coefficients were estimated via Hamiltonian Monte Carlo sampling, using weakly informative priors on all fixed effects. Posterior summaries are reported as means with 95% highest density intervals (HDI). All ANOVA tests were performed using the `statsmodels` Python package (Seabold & Perktold, 2010).

### Quantifying invariance across vessels

To identify which quantity participants held invariant across vessels, we computed, for each candidate measure and each participant, the proportion of that participant’s total variance attributable to the vessel. For a given participant, we partitioned the total sum of squares of the trial-level measurements about the participant’s mean into a between-vessel component and a residual trial-to-trial component:

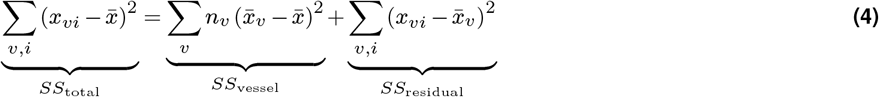

where *x*_*vi*_ is the measurement on trial *i* in vessel *v, n*_*v*_ the number of trials in vessel *v*, 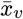 the participant’s mean in vessel *v*, and 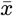 the participant’s overall mean. The proportion of variance due to vessel was then *SS*_vessel_*/SS*_total_, computed separately for each participant. A measure that a participant actively regulates should remain stable across vessels and thus produce a low value, whereas a measure that is affected by vessel geometry should yield a high value. We report the distribution of this proportion across participants for each of the six candidate measures (Fig. 2c).

To test whether the differences between measures were reliable, we computed bootstrap confidence intervals over participants. For each of 10,000 resamples, we drew 19 participants with replacement, computed the median proportion of variance due to vessel for each measure, and took the difference from headroom. We report the median difference and 95% bootstrap confidence interval for each comparison.

### Bootstrap analysis of variability

To compare within- and between-participant variability, we used a bootstrap procedure with 20,000 resamples. In each resample, we drew 20 trials without replacement from the relevant pool and computed the standard deviation of the headroom. For the within-versus between-participant comparison (Fig. 3b), we first uniformly sampled a participant, then drew 20 trials from that participant’s data (across tasks, containers, and vessels) to obtain the within-participant variability, and separately drew 20 trials from the remaining participants’ pooled data to obtain the between-participant variability. For the comparison across task conditions (Fig. 3d), we pooled all trials across participants, containers, and vessels, separately for the self-paced and fast conditions, and drew 20 trials from each pool. The resulting distributions are visualised as kernel density estimates.

### Data-Driven Dynamical System identification

We characterised the pouring dynamics using a data-driven dynamical systems framework. Specifically, we used SINDYc (Brunton et al., 2016b), a variant of the Sparse Identification of Nonlinear Dynamics algorithm that incorporates external control inputs into the state-space estimation. The system state at time *t* was defined as 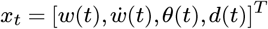, where *w*(*t*) is the liquid weight, 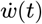 is the fill rate, *θ*(*t*) is the container rotation, and *d*(*t*) is the headroom to the rim of the vessel. The system was driven by the control input 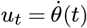, the container’s rotation rate.

#### Data Preprocessing for System Identification

To synchronize the data streams, scale data (*w*(*t*)) were downsampled from 1600 Hz to 120 Hz to match the motion capture data (*θ*(*t*)), resulting in a time step (*dt*) of 1*/*120 s. Both data streams were then filtered using a 5 Hz low-pass Butterworth filter. To estimate the fill rate 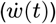 and rotation rate 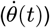—representing the first derivatives of fill weight and rotation, respectively—we applied a Savitzky–Golay filter (third-order polynomial, window size of 7) to compute smoothed finite differences. All container poses were transformed from the global coordinate system into the local reference frame of the active receiving vessel to maintain consistency across trials and participants.

#### System Identification

We estimated the pouring dynamics independently for each container rather than for specific container–vessel pairs, as the fluid flow-rate depends only on the container’s geometrical profile and is agnostic to the vessel type. Consequently, SINDYc system identification was performed on the state subspace 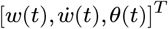 using 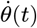 as the control input, and the vessel headroom (*d*(*t*)) was subsequently appended using the corresponding geometric solutions.

For each of the three containers, the dataset was split into an 80% training set and a 20% test set. The SINDYc basis function library was configured with second-order polynomials and a Fourier library composed of sine and cosine functions. The affine term was omitted, the Fourier basis was restricted to rotation metrics, and cross-term interactions were permitted. To ensure physical consistency, the predicted fill rate 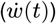 was wrapped in a ReLU function to prevent negative flow values. The resulting dynamics follow the state-space structure:

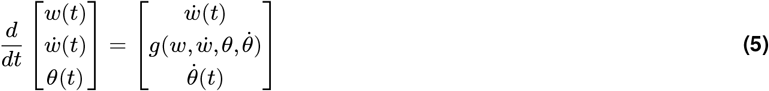

where *g*(·) represents the specific nonlinear acceleration of the fluid mass identified for each container:

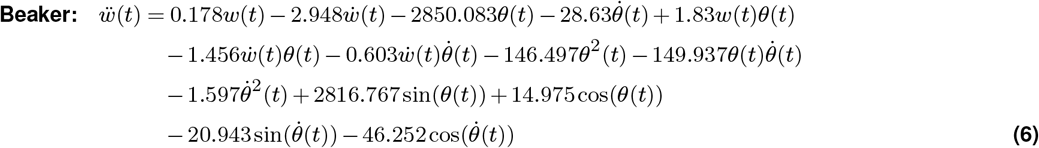

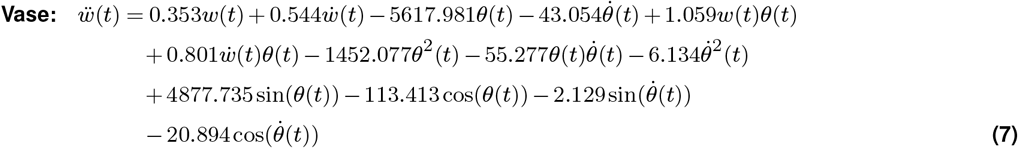

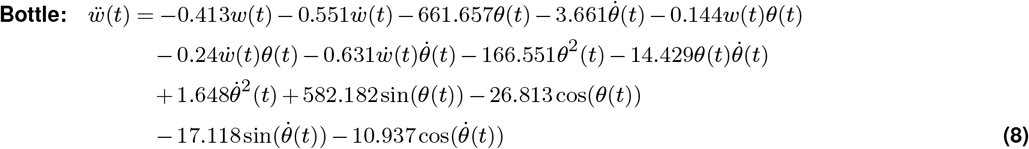

We evaluated the identified dynamics by comparing ground-truth trajectories against forward simulations using the corresponding ground-truth control inputs on held-out test trials. The resulting root mean square errors (RMSE) across all state variables for the Beaker, Vase and Bottle were 15.31, 23.65 and 19.24, respectively.

### Optimal Feedback Control Model

To solve the nonlinear control problem, we employed iterative Linear-Quadratic Gaussian control (iLQG; (Li & Todorov, 2007; Straub et al., 2023; Todorov & Li, 2005)). This method works by linearising the dynamics and approximating the cost function as locally quadratic around a nominal trajectory and computing the optimal linear control law,

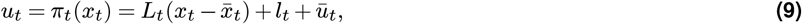

where *L*_*t*_ is the feedback gain and *l*_*t*_ is the affine control offset, both obtained from the backward recursion for the current nominal trajectory. The resulting policy is then applied in a forward rollout to generate an updated trajectory, and the procedure is repeated until convergence. Because our system involves partial observability, the controller is combined with an Extended Kalman Filter (EKF) acting as the optimal state estimator to track the internal belief states.

The state space was defined as 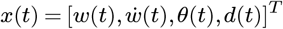 , with dynamics governed by the SINDYc-identified system described in Equation 5. The control input 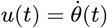 was corrupted by two forms of motor noise to reflect known properties of human motor control:

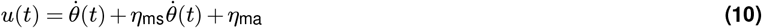

where 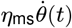 is a signal-dependent noise term, capturing the scaling of motor variability with signal magnitude (Harris & Wolpert, 1998), and *η*_ma_ is an additive noise term accounting for variability arising from the remaining degrees of freedom of the pouring action not captured by the single rotational control dimension.

The task was formalised under partial observability, assuming that the agent cannot directly perceive the internal fluid states (*w*(*t*), 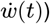 and observes only the headroom and the container rotation angle. The observation vector *y*(*t*) was therefore defined as:

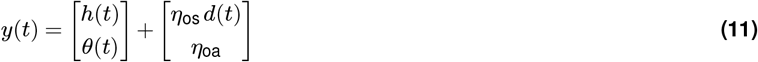

Observation noise on the headroom was modeled as signal-dependent *η*_os_, scaled by (*d*(*t*)), reflecting the empirical observation that variability in final fill level is higher at higher fill levels and vice versa. The container rotation state was corrupted by additive noise *η*_oa_, representing the agents’ proprioceptive uncertainty in container orientation. Given these definitions, the cost function used to derive the optimal control policy is given by Equation 1. We fixed *c*_task_ = 20 and *c*_rest_ = 2 across all analyses.

#### Noise parameters

The noise parameters were set separately depending on the purpose of each analysis. For the forward simulations illustrating the model’s ability to match variability in human trajectories (Fig. 4c), we used *η*_ms_ = 0.8, *η*_ma_ = 0.4, *η*_os_ = 0.1 and *η*_oa_ = 0.01. For the inverse modelling (Fig. 5), noise parameters were set to small values *η*_ms_ = *η*_ma_ = 10^−3^, *η*_os_ = *η*_oa_ = 10^−6^ so that the optimisation recovered cost function parameters from the mean trajectory shape rather than from the noise structure.

#### Signal-dependent observation noise analysis

To test whether signal-dependent observation noise is necessary to reproduce the observed relationship between mean headroom and its variability (Fig. 4d), we simulated the optimal controller across a range of fill levels and durations for each container–vessel combination. Specifically, we sampled 20 target fill levels uniformly spanning the range observed in the data for that combination, and for each fill level simulated the controller at five different time horizons drawn from the mean and standard deviation of the empirical fill times.

The noise parameters were determined per vessel in a two-stage procedure. First, we varied the signal-dependent motor noise *η*_ms_, additive motor noise *η*_ma_ and additive observation noise on rotation *η*_oa_, and manually selected the combination that produced variability levels closest to those observed in the data. Second, given these values, we swept the signal-dependent observation noise *η*_os_ across 17 uniform values between 0.1 and 2.0, and manually selected the value that best reproduced the observed relationship between mean headroom and its standard deviation, averaged across the five fill times and 20 fill levels. The resulting per-vessel noise parameters are reported in Table 1. For the comparison condition without signal-dependent observation noise, *η*_os_ was replaced by a small constant (10^−3^) while all other parameters were held fixed.

**Table 1.**
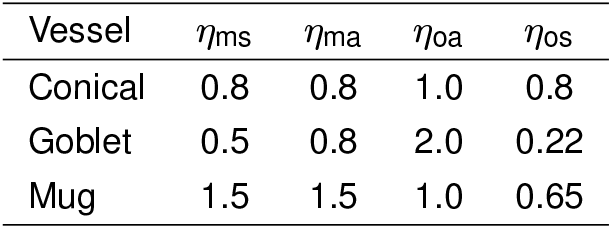
Noise parameters used in the signal-dependent observation noise analysis.

### Inverse modelling

Given the identified dynamics, control framework and cost function, we performed inverse modelling to estimate the cost function parameters from each participant’s trajectory data using nonlinear least-squares optimisation. This allowed us to interpret the observed behaviour and evaluate how control strategies varied across conditions, containers and vessels.

The cost function for pouring defined in Equation 1 contains five coefficients (*D*_tgt_, *c*_task_, *c*_eff_, *c*_flow_, *c*_rest_). Since *c*_task_ and *c*_eff_ jointly govern the trade-off between task success and motor effort, it suffices to fix one and optimise the other; we therefore fixed *c*_task_ = 20. We also fixed *c*_rest_ = 2, as this parameter only ensures that the container rotation *θ*(*t*) returns to zero after the pour and does not influence active pouring behaviour. We optimised over the remaining three free parameters *ϕ* = (*D*_tgt_, *c*_eff_, *c*_flow_) as they capture behaviourally relevant quantities: the energy expenditure (*c*_eff_), the degree to which flow-rate is regulated to avoid spillage (*c*_flow_), and the individual’s preferred fill level (*D*_tgt_). Furthermore, since iLQG is a fixed-horizon controller, the time horizon *T* was set to the duration of each individual trial during optimisation.

Optimisation was performed per participant, container, vessel, and task condition. Each dataset comprised up to 10 trials:

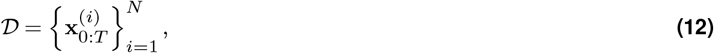

of which 80% were used for parameter estimation and the remaining 20% reserved for model evaluation and ΔBIC computation (Fig. 5d). Because the optimisation relies on trajectory fitting, we optimise only over the relevant subset of states x_sub_(*t*) = [*d*(*t*), *θ*(*t*)]^*T*^ . Tracking the fluid weight *w*(*t*) and fill rate 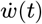 is redundant during optimisation as *d*(*t*) is directly computed from *w*(*t*) and 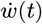 is the first derivative of *w*(*t*). Additionally, since the iLQG controller is stochastic, we simulated *M* = 50 trajectories given parameters *ϕ* and optimised over their mean trajectory,

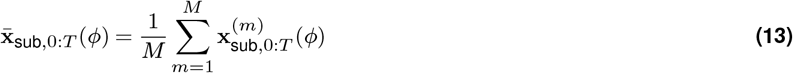

The optimal parameters 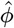 were estimated by minimizing the mean squared error between the simulated mean trajectory and the observed participant trajectories across all training trials using nonlinear least squares optimisation:

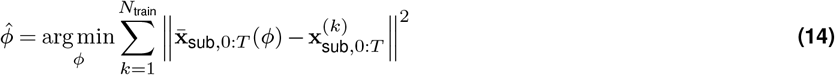

The objective was to find a single set of parameters per participant and condition that best explains the observed human behaviour. Before applying this pipeline to empirical data, we confirmed that the parameters can be uniquely recovered by testing the optimisation routine on synthetic data generated from known parameter values; validation details are provided in the Supplementary Material.

## Supplementary Information

### Cost function parameter analysis

To investigate the influence of each cost function parameter on the resulting control policies and state trajectories, we systematically varied each parameter independently while holding all others fixed, using the Mug vessel and Beaker container combination as reference environment. Since only the relative tradeoff between *c*_task_ and *c*_eff_ matters, rather than their individual values, we fix *c*_task_ and vary the remaining four parameters: *c*_eff_, *c*_flow_, *D*_tgt_, and *c*_rest_. We include *c*_rest_ here to show that it primarily affects the end phase of the pouring motion where the container resets to its initial rotation. For each parameter, we selected 10 values over a range (in either linear or log space depending on the parameter), fixing all others to the default values listed in Table 2. All simulations were run with a fixed time horizon of *T* = 650 timesteps, corresponding to 5.4 seconds.

**Table 2.** Default parameter values used in the cost function parameter analysis. When a given parameter is being varied, its default value is not used.

| Parameter | Default Value |
| --- | --- |
| $c_{\text{task}}$ | 20 |
| $c_{\text{eff}}$ | 0.05 |
| $c_{\text{flow}}$ | $1.51 \times 10^{-6}$ |
| $c_{\text{rest}}$ | 2.0 |
| $D_{\text{tgt}}$ | $0.2 H_v$ |

**Figure S1.**
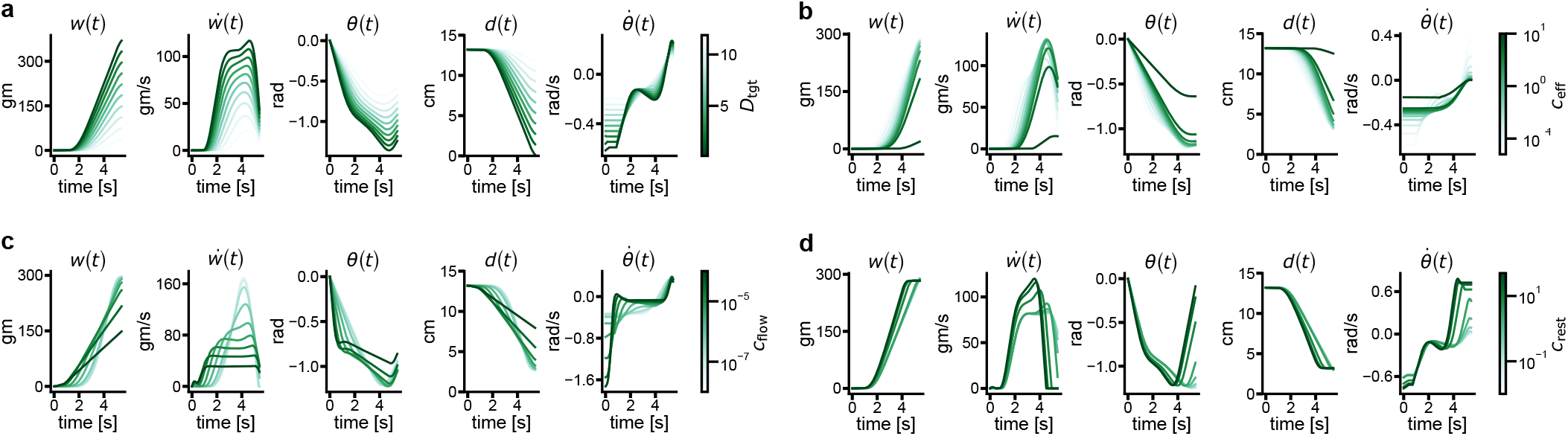
Effects of cost parameters on resulting trajectories

The results are shown in Figure S1. All four parameters affect the resulting trajectories in distinct and interpretable ways. In panel (a), varying *D*_tgt_ shows that the fill trajectory *d*(*t*) closely tracks the specified target, confirming that this parameter directly controls the agent’s stopping criterion. Panel (b) shows the effect of *c*_eff_ where at high values, 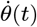 remains close to zero as the agent minimizes energy expenditure at the cost of not reaching the fill target, while at lower values the target is reached progressively earlier at the expense of higher energy expenditure. Panel (c) reveals how *c*_flow_ regulates the fill rate pattern: at low values, the flow profile takes a narrow parabolic shape with a high peak fill rate, whereas at high values the flow is suppressed into a step function pattern that stabilizes at a low, sustained rate. Finally, panel (d) demonstrates the role of *c*_rest_: at low values the container does not return to its resting orientation after filling completes, confirming that this term is a necessary component of the cost function, while at high values the rotation returns to zero as rapidly as possible.

### Inverse modelling validation

To verify that the IOC pipeline reliably recovers known parameters before applying it to human data, we first validated it on simulated data where ground truth values are known. For each task and all combinations of containers and vessels, we uniformly sampled 100 parameter sets for each of the three free parameters. The target headroom was sampled by drawing a fill fraction uniformly from [0.2, 1.0] and converting it to the corresponding distance to rim; the effort and flow coefficients were sampled from *c*_eff_ ∈ [10^−3^, 10^−1^] and *c*_flow_ ∈ [10^−7^ , 10^−5^]. For each sampled parameter set, we simulated trajectories using the iLQG-EKF controller setting the time horizon to the average time for the given container-vessel combination and then applied the inverse modelling procedure defined in Equation 14 to recover the parameters from the simulated data.

**Figure S2.**
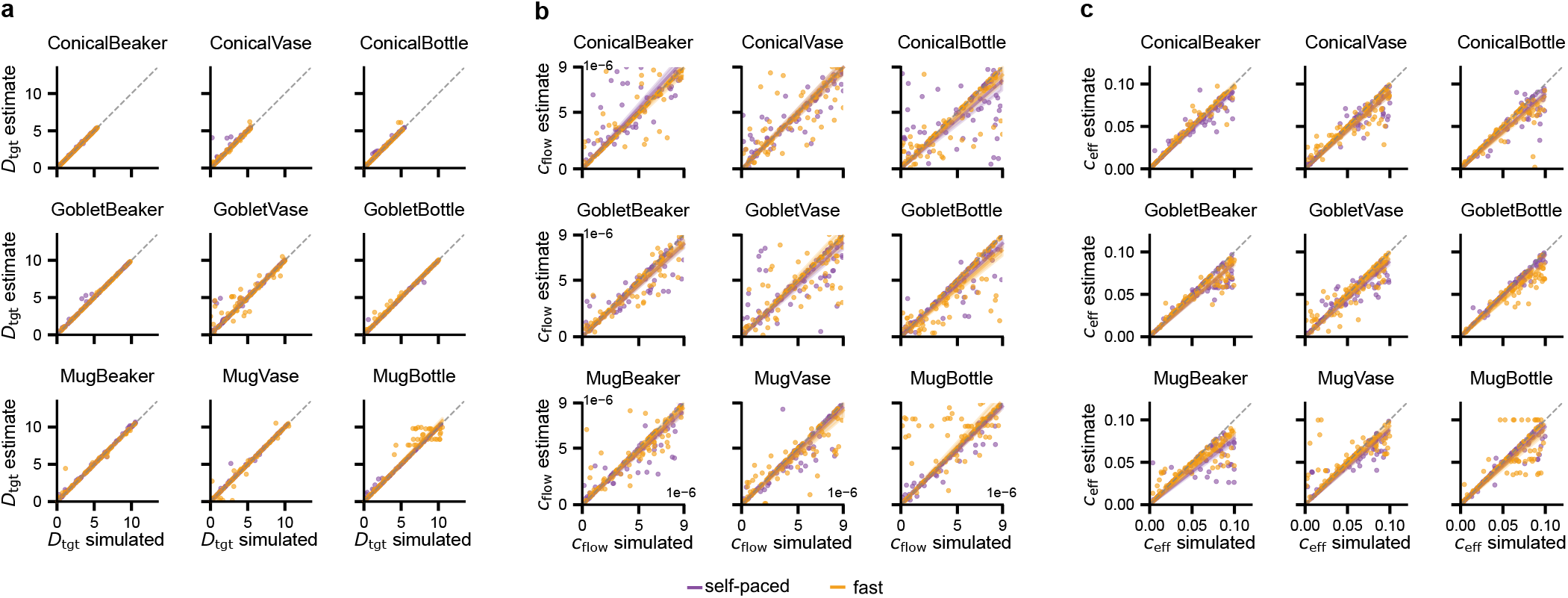
Recovered estimates versus ground-truth values for the three cost function parameters, across all container–vessel combinations. Dashed line indicates perfect recovery.

The results are shown in Figure S2, where the recovered estimates closely agree with the ground truth values across all parameters and conditions, confirming that the inference pipeline is identifiable and well-posed.

### Inferred trajectories under model ablations

To better understand the ΔBIC results reported in the main text, we show example trajectory fits from two representative participants for each of the three model variants (full model, no *c*_flow_, no *c*_task_) across three container-vessel combinations that span all containers and vessels in the experiment: Conical+Bottle, Goblet+Vase, and Mug+Beaker. Note that ablating *c*_eff_ is not considered here, as removing the effort cost causes the optimisation to fail entirely. The results are shown in Figure S3. The full model (red) closely tracks the ground truth data (black) across all conditions. Removing *c*_task_ (orange) causes the controller to remain idle throughout the trial, as without a task incentive there is no drive to pour. Removing *c*_flow_ (blue) eliminates the check on flow-rate, resulting in parabolic and physically implausible rotation profiles that would inevitably lead to spillage in practice.

**Figure S3.**
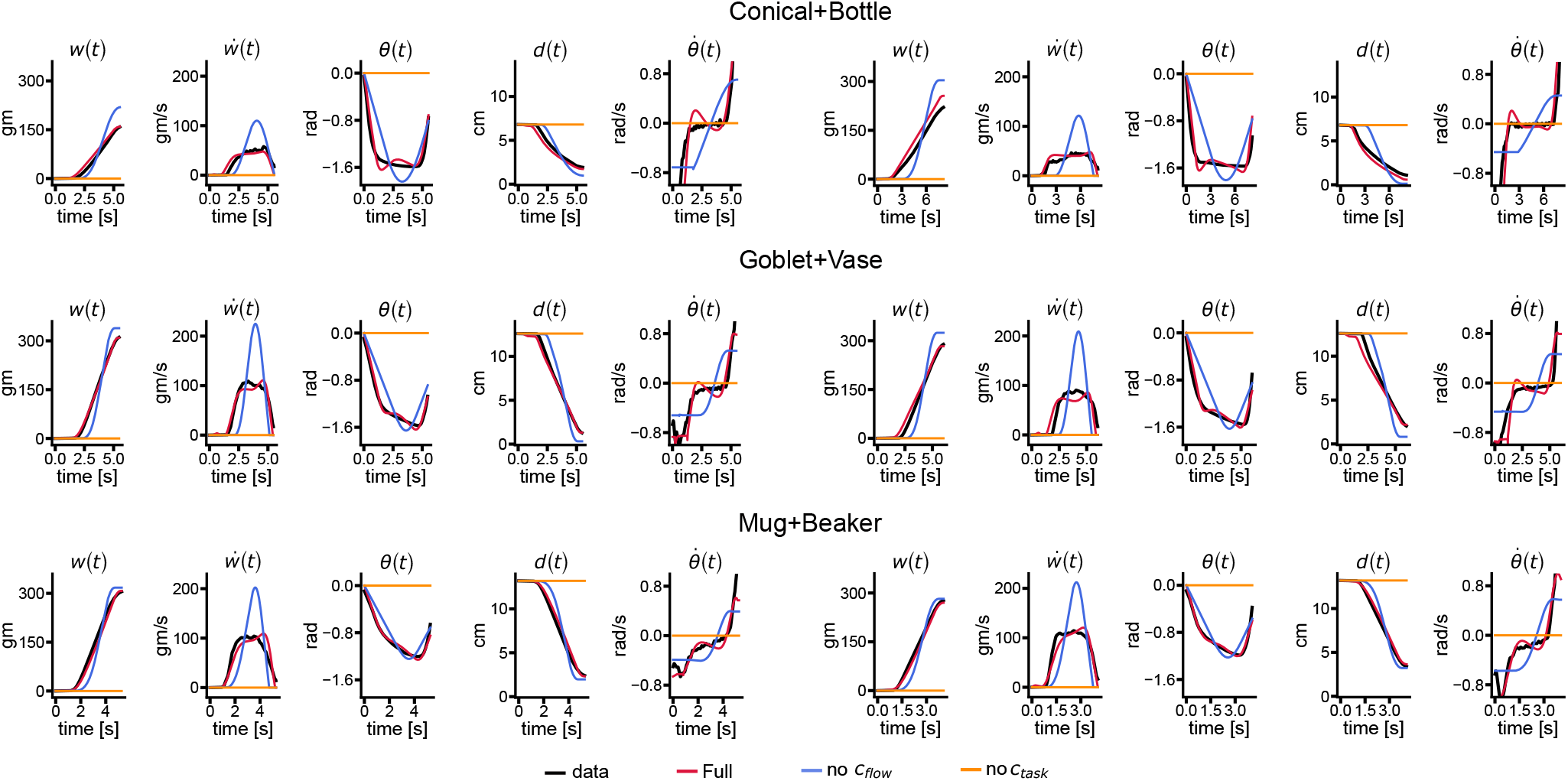
Observed data (black) and fits of the full model (red), without *c*_flow_ (blue), and without *c*_task_ (orange), for three representative container–vessel combinations.

## Notes

### Competing Interest Statement

The authors have declared no competing interest.

